# Outcome of TMS-based motor mapping depends on TMS current direction

**DOI:** 10.1101/371997

**Authors:** Jord JT Vink, Petar I Petrov, Stefano Mandija, Rick M Dijkhuizen, Sebastiaan FW Neggers

## Abstract

Navigated transcranial magnetic stimulation (TMS) in combination with electromyography (EMG) recordings can be used to map the brain regions in which TMS evokes motor-evoked potentials (MEPs) in certain muscles. Navigated TMS (nTMS) is used increasingly to identify the functional motor area of different muscles for clinical applications, including neurosurgical planning. However, the accuracy of TMS-based mapping of functional motor areas may depend on the TMS-induced current direction due to anisotropic cortical morphology, complicating association of the functional motor maps with neuroanatomical structures. Furthermore, it is not clear how well nTMS can distinguish nearby muscle representations on the cortical surface. We therefore investigated the functional motor maps obtained with posterior-to-anterior (PA) and lateral-to-medial (LM) TMS-induced currents within a spatially defined area by stimulating targets in a grid of locations over the left primary motor cortex in 8 healthy participants. Results were compared to functional MRI (fMRI) activation maps obtained using a voluntary opposing thumb movement task. We found that TMS applied with PA-induced currents identifies a motor area that is located significantly more anterior (8.7 – 10.4 mm depending on the muscle) with respect to an MEP motor area identified using LM-induced currents for the same muscle. Motor maps obtained with LM-induced currents show more overlap with the motor map identified using fMRI compared to PA-induced currents. In conclusion, the spatial representation of the MEP motor map identified by TMS is dependent on the direction of the induced current. These findings suggest that the application of nTMS using an LM-induced current direction corresponds best with the hand motor area as measured with fMRI.

## INTRODUCTION

Transcranial magnetic stimulation (TMS) is an increasingly popular non-invasive tool for diagnostic and therapeutic applications, which can be used to excite neurons in the brain through electromagnetic induction[1]. For diagnostic purposes, single pulses of TMS can be used to evoke activity in the primary motor cortex (M1), among other regions. If the current induced by TMS (TMS-induced current) exceeds a certain threshold, the evoked activity propagates through the cortico-spinal tract to the muscles, where it induces motor-evoked potentials (MEP). These MEPs can be recorded using electromyography (EMG)[2]. The waveform of the MEP can be quantified in terms of amplitude and latency. These electrophysiological measures reflect the excitability of the cortical neuronal populations and the total conduction time of the descending neuronal tracts, respectively.

TMS is frequently used in combination with MRI-guided neuronavigation, which allows mapping of the stimulated neuroanatomical targets that evoke responses in specific muscles.[3]. Using a standardized grid of stimulation targets, the functional motor areas of different muscles can be delineated systematically, based on the amplitudes of the MEPs as a function of stimulation location. In this way, TMS has been shown to be able to identify significantly separated motor area centers for different muscles in the upper extremity[4]. Neuronavigated TMS (nTMS) can potentially be of value in the preparation of neurosurgical interventions in the vicinity of the primary motor cortex, or to monitor changes in the primary motor cortex during the course of rehabilitation (of motor function) after stroke[5]–[11].

The MEP motor maps obtained with nTMS have been compared to anatomical MRI and functional MRI representations of the thumb area in the motor cortex[12]–[15]. These studies found that largest TMS-induced MEP is elicited anterior to the anatomical and functional representation of the thumb as measured with fMRI, with a discrepancy of up to 10mm. The majority of these studies used a posterior-to-anterior (PA) TMS-induced current (PA-induced current). In contrast to the other studies, Herwig et al. used a lateral-to-medial (LM) TMS-induced current (LM-induced current), roughly parallel to the precentral gyrus, and found an average anterior separation of 7.5mm[13]. It was speculated that the discrepancy was caused by the orientation of the TMS coil (or the physical properties of the TMS coil). However, it was not systematically investigated whether this discrepancy could arise from the direction of the TMS-induced current with respect to the cortical morphology.

The widespread use of nTMS in the identification of the functional motor area for clinical purposes is restricted due to the limited understanding of the interaction of the TMS-induced current and the neuronal populations in the brain. For example, TMS effects have been shown to depend on the orientation of the TMS coil, and thus the direction of the induced current, with respect to the local morphology of the stimulated area. The orientation of the TMS-induced current with respect to the underlying motor cortex has been shown to affect the amplitude of the MEPs, with induced currents perpendicular to the orientation of the sulcal wall (a PA current direction) of the precentral gyrus resulting in larger MEP amplitudes[16][17]. This has also been observed for stimulation of the visual cortex through the induction of phosphenes (visual illusions evoked by TMS)[18]. Others have found that the latency of MEPs depends on the induced current direction and concluded that LM-induced currents preferentially evoke activity in other cortical elements than PA-induced currents[19]–[21].

Based on the aforementioned work, we speculate that the spatial distribution of cortical targets where MEPs are elicited varies as a function of the orientation of the current induced by TMS. Understanding the effect of the TMS-induced current direction on the spatial distribution of MEP-eliciting cortical targets (from now on referred to as the MEP motor map) is crucial for the interpretation of functional motor maps obtained with nTMS. This is especially important as nTMS is used increasingly in the identification of the functional motor area in the planning of neurosurgical interventions[5]–[8][11][22].

We therefore systematically investigated whether PA and LM TMS-induced current directions give rise to different MEP motor maps. This was done by stimulating targets in a grid over the left primary motor cortex, using a novel nTMS-EMG technique, and by comparing the location of the center of gravity of MEP motor maps to the location of the maximum response to voluntary thumb movements, as measured with functional MRI in the same participants. In addition, we tested the ability of nTMS to distinguish MEP motor map centers for different muscle groups, as previously reported[4].

## 2. MATERIALS AND METHODS

The experimental procedure was approved by the medical ethical committee of the University Medical Center Utrecht (UMCU), Utrecht, The Netherlands. MRI data were acquired in 8 healthy right-handed participants (4 male; 4 female; mean age: 26.4). All participants provided written informed consent and were screened for MRI and TMS exclusion criteria. During the experimental procedure, we strictly adhered to the guidelines and recommendations for TMS endorsed by the International Federation for Clinical Neurophysiology[23].

### 2.1 Materials

All MR experiments were performed on a 3T MR scanner (Philips Achieva, Best, The Netherlands, www.philips.nl). Navigated TMS was performed using the Neural Navigator 3.1 ‘navigated MEP’ (Brain Science Tools BV, De Bilt, The Netherlands, www.brainsciencetools.com). A monophasic magnetic stimulator Neuro-MS (Neurosoft, Ivanovo, Russia, www.neurosoft.com) with a figure-of-8 TMS coil 100mm FEC-02-100 (Neurosoft, Ivanovo, Russia) and a 4 channel EMG amplifier Neuro-EMG (Neurosoft, Ivanovo, Russia) were used to evoke and record MEPs simultaneously. The EMG signals were recorded, amplified and digitized with a sampling frequency of 20 kHz. The Neurosoft software controlling the magnetic stimulator and EMG device was designed to communicate in real time with the Neural Navigator software, allowing real time capturing of the TMS coil position and orientation and visualization of MEPs on the cortical surface.

### 2.2 Methods

The experiment was divided into two parts: an MRI session and a TMS-EMG session (Fig. 1). During the MRI session, a T1 weighted anatomical scan and a functional MRI time series were obtained. The first scan was used for neuronavigation during the TMS-EMG session, while the latter was used to identify the maximum blood-oxygen-level-dependent (BOLD) response in response to voluntary thumb movements. In the TMS-EMG sessions, the same target points were stimulated using a PA- and LM-induced current, separately.

**Figure 1.**
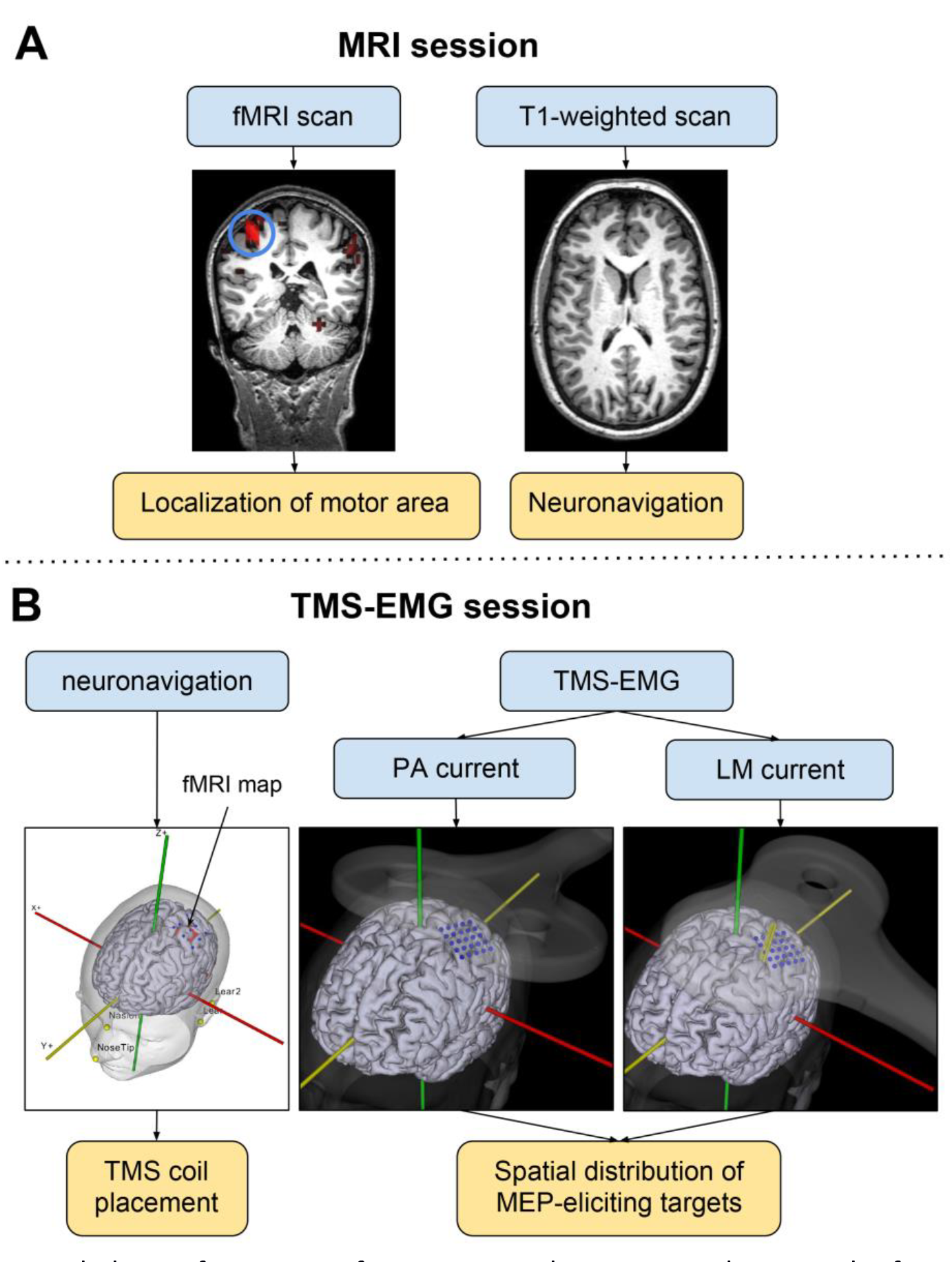
Schematic overview of the experimental procedure. **Panel A.** The images acquired during the MRI session. **Panel B.** The 2 subsequent TMS-EMG sessions in which different TMS-induced current directions were used. The yellow boxes below the figures denote the most important derived measures from each procedure. PA current: Posterior-to-anterior induced current. LM current: lateral-to-medial induced current.

#### 2.2.1 MRI session

First, a T1-weighted anatomical scan was acquired with a TR/TE of 10.015/4.61ms, a flip angle of 8°, voxel size of 0.75x0.75x0.8mm^3^, scan duration of 677s, and 225 slices with a slice gap of 0mm.

Thereafter, a single-shot EPI sequence was acquired with 250 dynamics, a TR/TE of 2,000/23ms, flip angle of 70°, voxel size of 4x4x4mm^3^, scan duration of 510s, and 30 slices with a slice thickness of 3.6mm and a slice gap of 0.4mm. During the EPI sequence, the participant was instructed to perform opposing thumb movements upon presentation of an auditory cue (which was manually presented every 10 to 16 s to avoid habituation). The movements of the right abductor pollicis brevis (APB) muscle were recorded using a wireless MR-compatible electrocardiography device (Invivo, Best, The Netherlands). EMG was recorded at a frequency of 496Hz, in order to detect voluntarily-induced muscle activity, allowing us to reconstruct the timing of the voluntary thumb movements.

#### 2.2.2 TMS-EMG session

For each participant, 3D surface renderings of the skin and cortex were obtained from the T1-weighted MRI and visualized in the Neural Navigator software. Subsequently, the location of the hand area in the left M1 was derived from the statistical activation map computed from the BOLD data of the fMRI scan (see section 2.4) and marked in the Neural Navigator. In the Neural Navigator software, a 5 by 5 grid of stimulation targets with 7 mm distance between the targets and a total area of 2.8 by 2.8 cm was centered on the hand area in the left M1. This grid was placed to aid the real time targeting of the TMS pulse on preplanned locations on the motor cortex.

The participant was seated comfortably on a chair with the arm resting on a table. Surface electrodes were placed over the right first dorsal interosseous (FDI), abductor digiti minimi (ADM) and extensor carpi radialis (ECR) muscles in a belly-tendon montage and the ground electrode was attached to the left wrist. The navigation procedure was performed using at least 6 facial landmarks to assure accurate alignment between tracker space and MRI space. Head movement compensation was performed by the neural navigator in real time, using continuous readings of two position tracking sensors attached to the head. This allows to spatially correct for small head movements in real time, and to perform accurate neuronavigation throughout the procedure[15].

The resting motor threshold (RMT) determination and single pulse stimulation of the grid targets was performed for the two TMS-induced current directions (PA and LM current directions), separately. A PA current was induced using a PA TMS coil orientation with the coil handle oriented perpendicular to the orientation of the individual participant’s precentral gyrus (at a 20° to 40° angle with the mid-sagittal plane depending on individual cortical morphology)(Fig. 1). An LM current was induced using a LM TMS coil orientation with the coil handle oriented parallel to the orientation of the precentral gyrus (at a 90° to 110° angle with the mid-sagittal plane depending on individual cortical morphology)(Fig. 1). The RMT was determined at the location where the maximum MEP was observed by probing different target in the vicinity of the precentral gyrus for the PA- and LM-induced current directions, separately. This method was chosen as from previous literature it is known that a higher RMT is needed for LM orientation, in order to avoid getting insufficient MEP responses for the LM session[16][17]. The RMT was obtained by increasing the stimulator output until MEPs with a peak-to-peak amplitude of over 50 μV were observed in the EMG recording of the contralateral FDI muscle in 5 out of 10 trials with an interstimulus interval of 7s[2][23]. For each session, the machine output was then set at 120% of the previously determined RMT. Targets in the grid were stimulated with 5 consecutive TMS pulses with an interstimulus interval of 3s. A total of 125 TMS pulses was delivered per TMS current direction (total of 250 pulses). The targets were stimulated in a random order to make sure that the stimulation order would not affect statistical analysis. The target grids were obtained for both TMS-induced current directions (PA and LM), separately.

The EMG trace and TMS coil position, orientation and angulation (with respect to the head) were captured during TMS pulse delivery (Fig. 2). In combination with head movement compensation, this allowed accurate registration of stimulated cortical targets. The data were visualized in the Neural Navigator 3.1 in real time and exported to an XML format text file for further analysis in Matlab.

**Figure 2.**
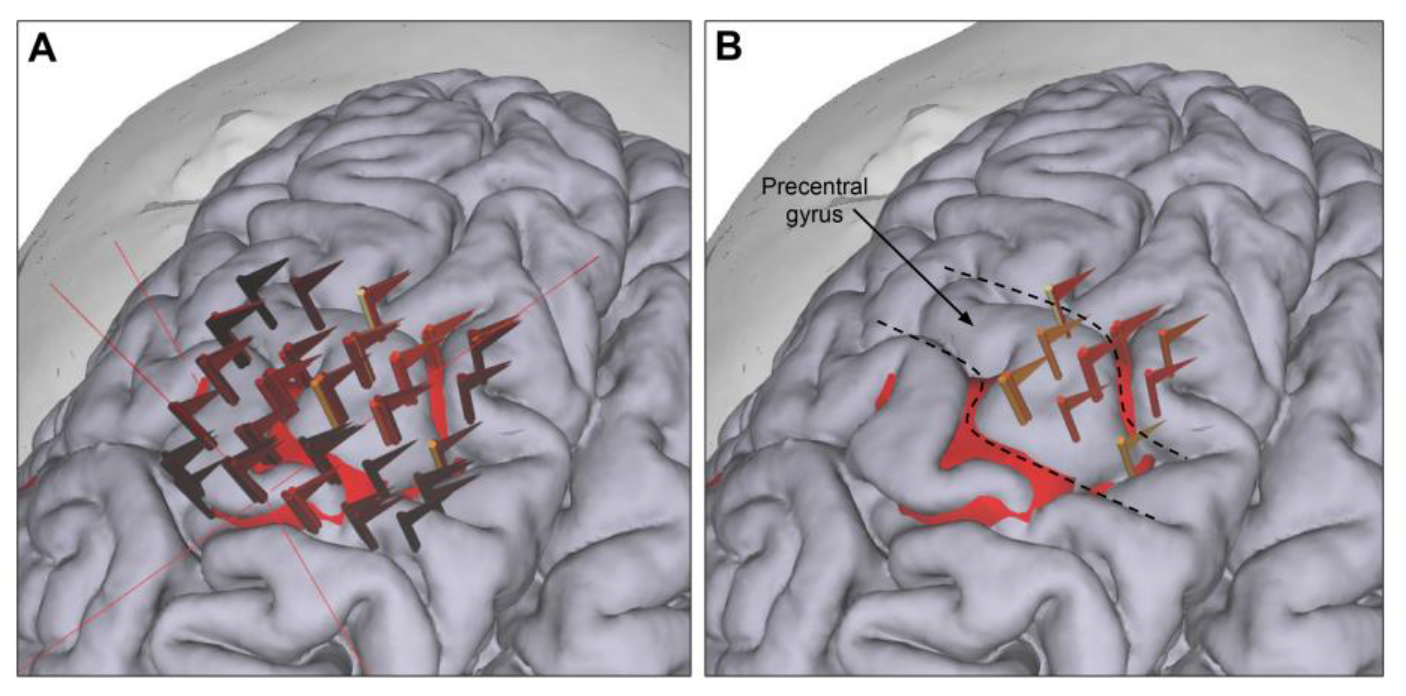
The TMS coil locations and orientations were captured during TMS pulse delivery. The flags placed at intersection of the TMS coil isocenter and the cortical surface upon TMS pulse delivery. The flag points in the direction of the TMS-induced current and the color coding indicates the MEP amplitude (0 – 3 mV). The red surface shows the fMRI-based hand area. **Panel A.** All stimulated targets. **Panel B.** All stimulated targets with an MEP amplitude of more than 1 mV.

#### 2.2.3 Validation tests

We retested the spatial accuracy of the navigation system (original tests are reported in Neggers et al 2004)[15]. A liquid probe was attached to a position on the scalp of one of the participants before acquisition of the T1-weighted MRI, so that the target position on the scalp could be visualized in the individual MR images, allowing validation of the accuracy of the neuronavigation system. Next, we used this MR image for neuronavigation, and navigated to the position of the liquid probe on the scalp visible in the surface rendering in the Neural Navigator. Finally, we repeatedly determined the distance between the position of the neuronavigation pointer and the liquid probe on the MR image. The distance was at most 3mm, which is within the reported accuracy of neuronavigation of 4mm[15].

We also determined whether the maximum B-field of the TMS coil was indeed located in the middle of the two coil windings, the TMS coil isocenter, which was used to navigate the TMS coil. This is important to establish as a deviation between the actual and the assumed isocenter would cause an error in the location of TMS-based MEP motor area with respect to the neuroanatomy. We investigated this by repeatedly discharging the TMS coil mounted rigidly at 1cm over a glass plate of iron filings and investigated the distribution of the filings afterwards. It is known that a small heap of filings piles up at the isocenter of a magnetic field where the field lines run exactly perpendicular to the surface. We found that this heap and hence the maximum B- and induced E-field was indeed located exactly at the theoretical TMS coil isocenter that we calibrated using The Neural Navigator. We could confirm, however, that the TMS coil we used was fully according to its specifications in this respect.

#### 2.2.4 Data analysis

All data were analyzed using custom scripts and SPM12[24] in a Matlab 2014a environment (Mathworks Inc., USA). For each participant, the T1 weighted image was segmented using unified segmentation in SPM12 to obtain a grey matter, white matter and CSF mask[25]. The location of the maximum response to thumb movements was obtained from the statistical activation map from the BOLD data of the fMRI scan (fMRI-based hand area). The statistical map was constructed based on a subject-level event-related general linear model (GLM) analysis, in which the thumb movements were modeled with the canonical hemodynamic response function (HRF). The timing of the thumb movements was obtained from the EMG recordings which were acquired during MRI acquisition using custom Matlab code. The GLM included two nuisance regressors: the average BOLD signal in the white matter and the CSF. Statistical images were constructed based on a T-statistic with the T-threshold at P < 0.05, family wise error (FWE) whole-brain corrected[24].

The EMG traces of all muscles were filtered with a low pass filter with a cutoff frequency of 500Hz, in order to eliminate the noise caused by the electromagnetic tracking device from the EMG signals. A custom peak detection algorithm (minima and maxima were determined between 15 and 40ms post-stimulus) was used to determine the peak-to-peak amplitude of the MEP. The 5 MEP amplitudes per target grid point were averaged together. Next, the center of gravity (COG) of the MEP motor map was calculated. This was done by weighing all the target coordinates by the amplitude of the corresponding MEP. MEPs with amplitude less than 50μV or MEPs with noise levels exceeding the maximum MEP amplitude were weighted zero. Separations between COGs of MEP motor maps obtained with different coil orientations or muscles were determined by calculating the absolute distance between the X and Y coordinates (anterior-posterior and lateral-medial direction, respectively) of the COG, since separation in the Z direction is predominantly dependent on brain morphology and sulcal depth, as all grids were only slightly tilted with respect to the axial plane.

The latencies of all the MEPs within the MEP motor maps obtained using PA and LM-induced current directions were also compared.

#### 2.2.5 Statistical analysis

The difference between the RMTs for PA and LM induced current directions was assessed through a one-sided Student’s paired t-test, because the literature clearly states that the RMT is higher for an LM induced current (parallel to the precentral gyrus)[26]. The separation in anterior-posterior direction between the COG for a PA TMS-induced current and the location (voxel) with the maximum BOLD response to voluntary thumb movements was also determined through a one-sided paired Student’s t-test, guided by existing evidence that the COG lies anterior of the maximum BOLD response[12]. All the other separations were evaluated with two-sided paired Student’s t-tests, as we had no a-priori hypotheses for these comparisons. Statistical tests evaluating separations in the X and Y direction were corrected for multiple comparisons by using a Bonferroni-corrected alpha of 0.025 (2 comparisons). Statistical tests between the cortical representations of different muscles were also corrected for multiple comparisons, resulting in a Bonferroni-corrected alpha of 0.0125 (4 comparisons). Finally, differences in latency were evaluated with a one-sided paired Student’s t-test because prior literature suggests increased latencies for a PA-induced current with respect to LM-induced current[19][27].

## 3. RESULTS

The RMT was determined separately for sessions with PA- and LM-induced currents. We found a significant increase in RMT (mean increase: 5.3%) for coil orientations with an LM-induced current with respect to a PA-induced current (P = 0.0005, T = 5.4, df = 7; Table 1).

**Table 1.** The resting motor thresholds (% of maximum stimulator output) obtained with PA and LM-induced currents for all participants. The values below the double line are the average RMTs.

| Participant number | RMT (FDI muscle) |  |
| --- | --- | --- |
|  | PA current | LM current |
| 1 | 37 | 41 |
| 2 | 40 | 41 |
| 3 | 31 | 33 |
| 4 | 38 | 44 |
| 5 | 40 | 48 |
| 6 | 41 | 50 |
| 7 | 41 | 48 |
| 8 | 27 | 35 |
|  | 36.9 | 42.5 |

FDI: first dorsal interosseous; PA: posterior-to-anterior; LM: lateral-to-medial.

The COGs of the MEP motor maps of the FDI, ADM and ECR muscles obtained using a PA-induced current were located significantly more anterior with respect to the COGs obtained with an LM-induced current, with average separations of 9.3, 10.4 and 8.7 mm respectively (Table 2). Figure 3 shows MEP motor maps of three different participants obtained with PA and LM TMS-induced current directions and figure 4 shows the relative distances between the COGs of the MEP motor maps obtained using PA-induced currents with respect to the COGs using LM-induced currents for all participants. The figures clearly show that the distribution of MEPs evoked with a PA-induced current is located anterior with respect to MEPs evoked with an LM-induced current.

**Figure 3.**
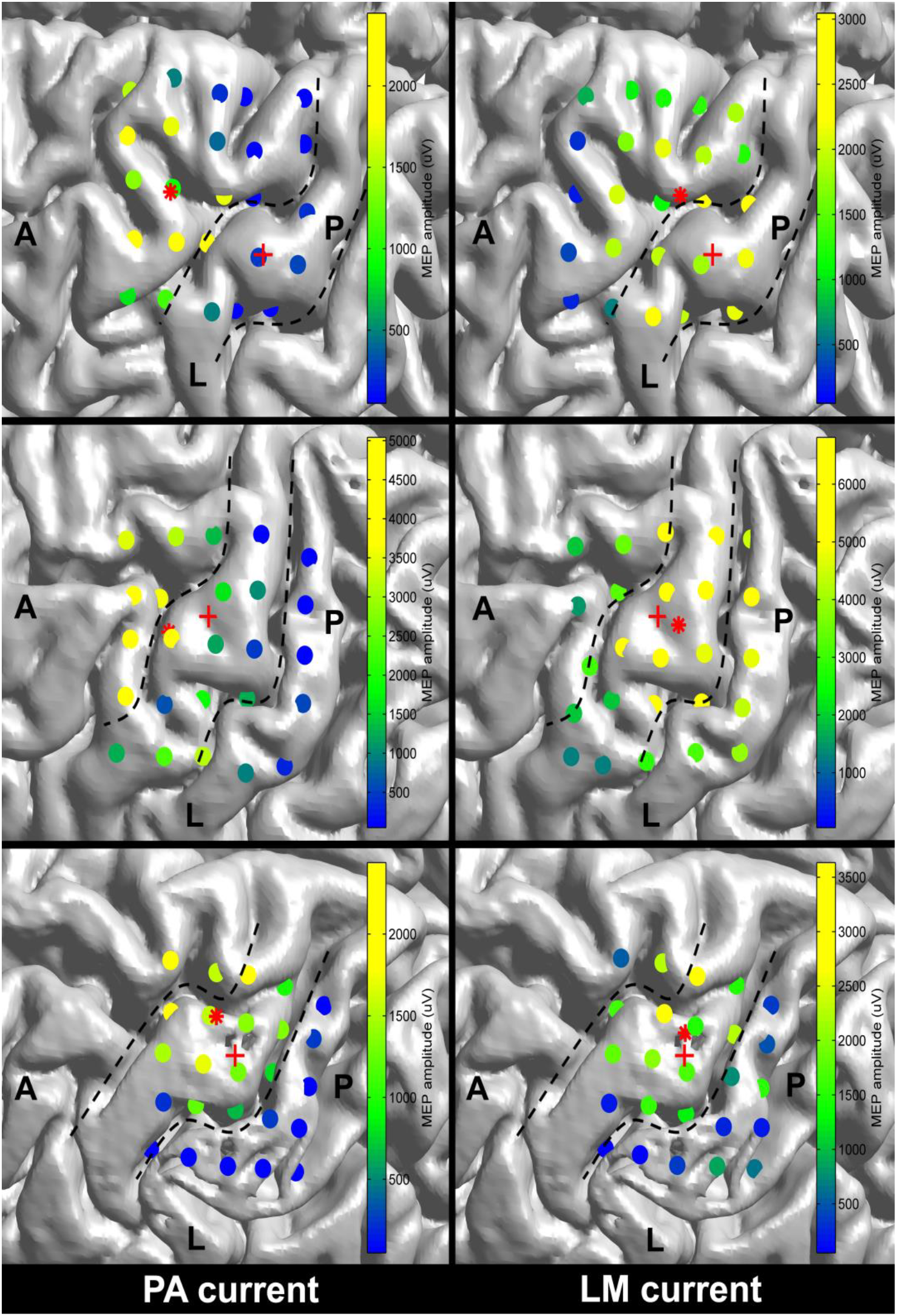
MEP motor maps of the FDI muscle for 3 different participants and PA- and LM-induced current directions projected on individual brain morphology. The color represents the amplitude of the MEP as indicated by the colorbar. The red plus sign (+) shows the location of the fMRI-based hand area and the red star sign (*) shows the location of the center of gravity of the MEP motor map. The dotted lines indicate the boundaries of the precentral gyrus. A: anterior; P: posterior; L: lateral; FDI: first dorsal interosseous.

**Figure 4.**
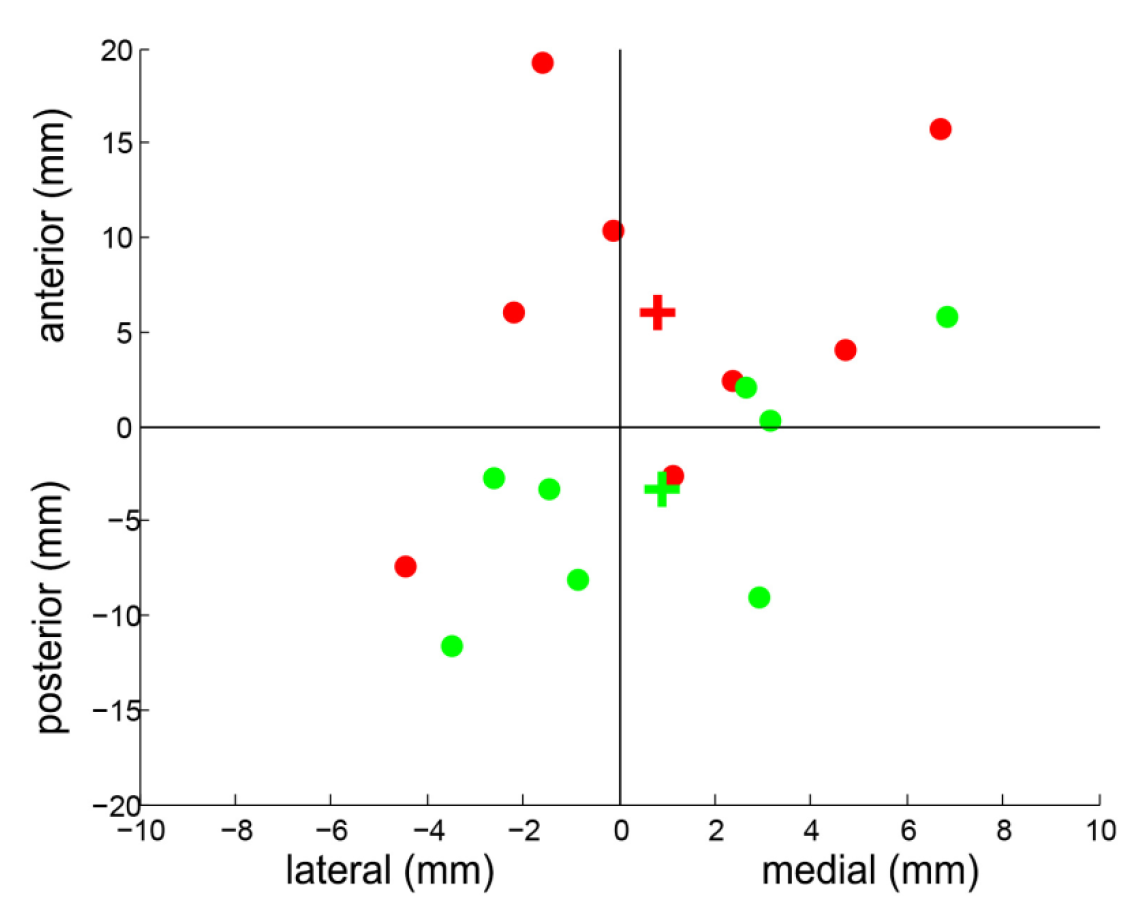
The relative locations of the COGs of MEP motor maps (FDI muscle) obtained with two different TMS-induced current directions with respect to the location of the maximum BOLD response to opposing thumb movements. Each dot represents an individual participant. The COGs obtained using PA- and LM-induced currents are shown as red and green dots, respectively. The mean COGs are indicated by a plus (+). It can be observed that PA-induced currents evoke anteriorly located COGs.

**Table 2.** Separations between the TMS-based COGs obtained with a PA-induced current with respect to an LM-induced current (df = 7). Significant correlations are indicated by an asterisk.

| Muscle | X-coordinate separation<br>(Anterior(+)/Posterior(-)) |  |  |  | Y-coordinate separation<br>(Medial(+)/Lateral(-)) |  |  |  |
| --- | --- | --- | --- | --- | --- | --- | --- | --- |
|  | Mean<br>(mm) | SD (mm) | T | P | Mean<br>(mm) | SD (mm) | T | P |
| FDI | 9.3 | 5.4 | 4.6 | 0.003* | -0.1 | 2.5 | -0.1 | 0.900 |
| ADM | 10.4 | 5.0 | 5.5 | <0.000* | -0.4 | 2.6 | -0.4 | 0.712 |
| ECR | 8.7 | 5.5 | 4.2 | 0.004* | -1.1 | 2.3 | -1.3 | 0.251 |

FDI: first dorsal interosseous; ADM: abductor digiti minimi; ECR: extensor carpi radialis.

The COGs of the MEP motor maps of the FDI, ADM and ECR muscles obtained with PA-induced currents were located anterior to the fMRI-based hand area, while a medial-lateral separation was negligible (Table 3). These separations were only significant for the ADM muscle with a separation of 6.0 mm, but showed a strong trend for the other muscles. The COGs of the MEP motor maps of the FDI, ADM and ECR muscles were not significantly separated from the fMRI-based hand area (Table 4).

**Table 3.** Separations between the TMS-based COGs obtained using a PA-induced current and the fMRI-based hand area (df = 7). Significant correlations are indicated by an asterisk.

| Muscle | X-coordinate separation<br>(Anterior(+)/Posterior(-)) |  |  |  | Y-coordinate separation<br>(Medial(+)/Lateral(-)) |  |  |  |
| --- | --- | --- | --- | --- | --- | --- | --- | --- |
|  | Mean<br>(mm) | SD (mm) | T | P | Mean<br>(mm) | SD (mm) | T | P |
| FDI | 6.0 | 8.4 | 1.9 | 0.050 | 0.8 | 3.5 | 0.6 | 0.566 |
| ADM | 6.0 | 8.2 | 1.9 | 0.047* | 2.2 | 3.2 | 1.7 | 0.121 |
| ECR | 4.8 | 8.0 | 1.6 | 0.080 | 1.1 | 3.3 | 0.9 | 0.410 |

FDI: first dorsal interosseous; ADM: abductor digiti minimi; ECR: extensor carpi radialis.

**Table 4.** Separations between the TMS-based COGs obtained using an LM-induced current and the fMRI-based hand area (df = 7). Significant correlations are indicated by an asterisk.

| Muscle | X-coordinate<br>(Anterior(+)/Posterior(-)) |  |  |  | Y-coordinate<br>(Medial(+)/Lateral(-)) |  |  |  |
| --- | --- | --- | --- | --- | --- | --- | --- | --- |
|  | Mean<br>(mm) | SD (mm) | T | P | Mean<br>(mm) | SD (mm) | T | P |
| FDI | -3.3 | 5.6 | -1.6 | 0.921 | 0.9 | 3.3 | 0.7 | 0.489 |
| ADM | -4.3 | 5.7 | -2.0 | 0.958 | 2.5 | 2.9 | 2.3 | 0.057 |
| ECR | -3.9 | 6.2 | -1.7 | 0.930 | 2.2 | 3.3 | 1.7 | 0.125 |

FDI: first dorsal interosseous; ADM: abductor digiti minimi; ECR: extensor carpi radialis.

The COGs of MEP motor maps of some of the individual muscles were significantly separated. The COG of the MEP motor map of the FDI muscle was significantly separated in the lateral direction with respect to the COG of the ADM muscle for both TMS current directions (Table 5; Table 6; Fig. 5; Fig. 6). Details on the separations between the COGs of MEP motor maps for different muscles can be found in tables 5 and 6.

Finally, the latencies of all the MEPs in the MEP motor area obtained with a PA- and LM-induced current are similar, with 23.5 and 23.6ms respectively (Table 7).

**Figure 5.**
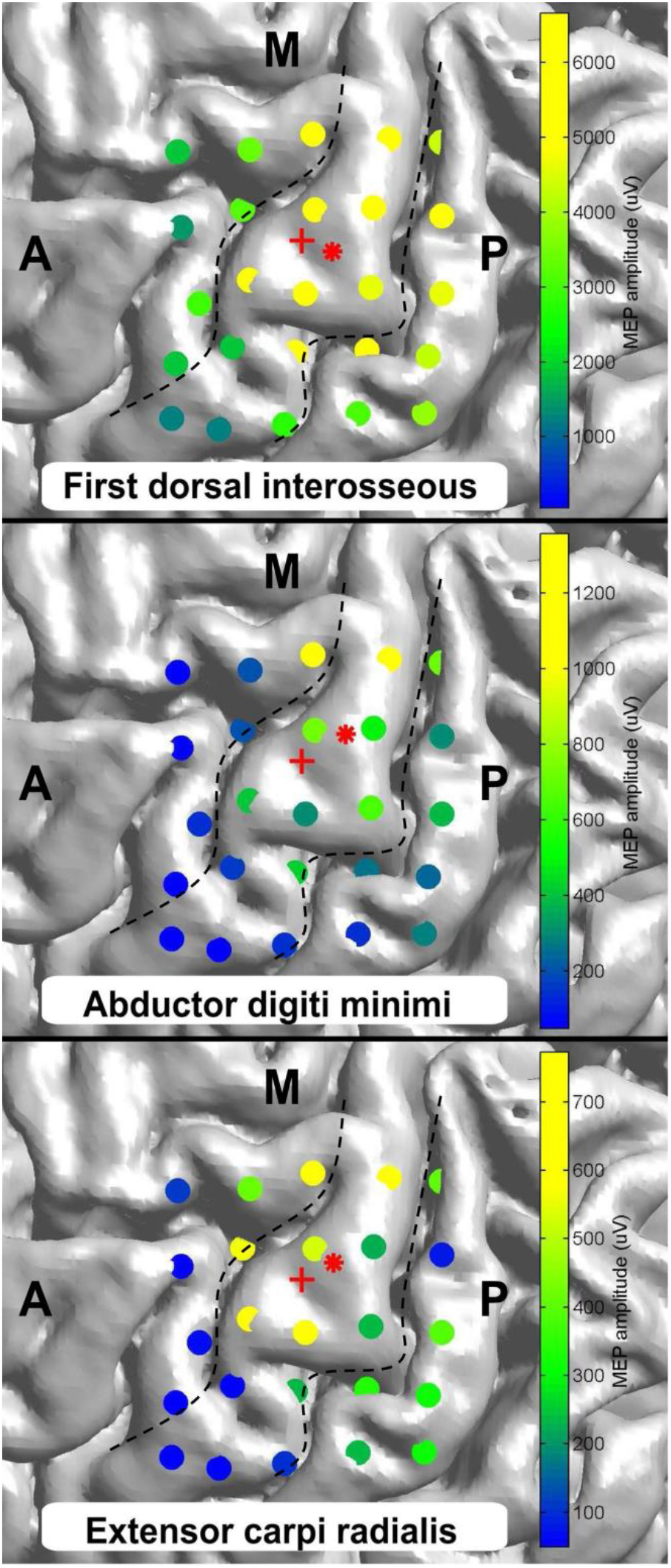
MEP motor maps of 3 different muscles obtained with an LM-induced current visualized on individual brain morphology of one representative participant. The color represents the amplitude of the MEPs as indicated by the colorbar. The red plus sign (+) shows the location of the maximum BOLD response to opposing thumb movements and the red star sign (*) shows the location of the center of gravity of the MEP motor map. The dotted lines indicate the boundaries of the precentral gyrus. A: anterior; P: posterior; M: medial.

**Figure 6.**
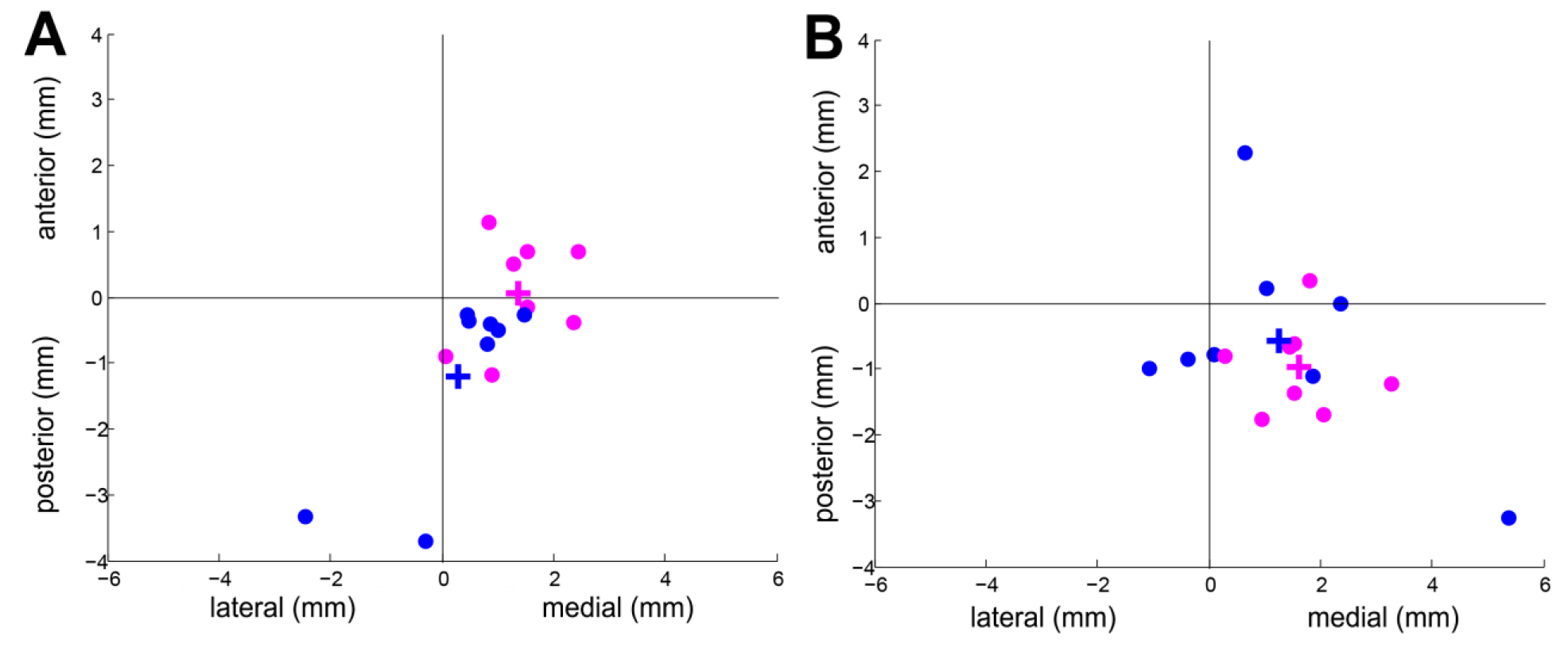
The relative locations of the COGs of the MEP motor maps of the ADM and ECR muscles with respect to the COG of the FDI muscle. The COG of the FDI muscle is significantly more lateral compared to the COG of the ADM muscle for both induced current directions. The COGs of the ADM and ECR are shown in pink and blue circles (o), respectively. Each dot represents an individual participant. The mean COGs are indicated by a plus (+). **Panel A.** COGs of MEP motor maps obtained with PA induced currents. **Panel B.** COGs of MEP motor maps obtained with LM induced currents.

**Table 5.** Separations between the TMS-based COGs of the investigated muscles obtained using a PA-induced current (df = 7). FDI-ADM indicates a separation of the FDI with respect to the ADM. Significant correlations are indicated by an asterisk.

| Muscles | X-coordinate separation<br>(Anterior(+)/Posterior(-)) |  |  |  | Y-coordinate<br>(Medial(+)/Lateral(-)) |  |  |  |
| --- | --- | --- | --- | --- | --- | --- | --- | --- |
|  | Mean<br>(mm) | SD (mm) | T | P | Mean<br>(mm) | SD (mm) | T | P |
| FDI-ADM | -0.1 | 0.8 | -0.2 | 0.865 | -1.4 | 0.8 | -4.9 | 0.002* |
| FDI-ECR | 1.2 | 1.4 | 2.3 | 0.052 | -0.3 | 1.2 | -0.7 | 0.505 |
| ADM-ECR | 1.2 | 0.9 | 3.8 | 0.007* | 1.1 | 0.9 | 3.2 | 0.016 |

FDI: first dorsal interosseous; ADM: abductor digiti minimi; ECR: extensor carpi radialis.

**Table 6.** Separations between the TMS-based COGs of the investigated muscles obtained using an LM-induced current (df = 7). FDI-ADM indicates a separation of the FDI with respect to the ADM. Significant correlations are indicated by an asterisk.

| Muscles | X-coordinate separation<br>(Anterior(+)/Posterior(-)) |  |  |  | Y-coordinate separation<br>(Medial(+)/Lateral(-)) |  |  |  |
| --- | --- | --- | --- | --- | --- | --- | --- | --- |
|  | Mean<br>(mm) | SD (mm) | T | P | Mean<br>(mm) | SD (mm) | T | P |
| FDI-ADM | 1.0 | 0.7 | 4.0 | 0.005* | -1.6 | 0.9 | -5.2 | 0.001* |
| FDI-ECR | 0.6 | 1.6 | 1.0 | 0.336 | -1.2 | 2.0 | -1.7 | 0.124 |
| ADM-ECR | -0.4 | 1.8 | -0.6 | 0.549 | 0.4 | 2.0 | 0.5 | 0.615 |

FDI: first dorsal interosseous; ADM: abductor digiti minimi; ECR: extensor carpi radialis.

**Table 7.** Difference in latency of the MEPs within the MEP motor map obtained with a PA-induced current with respect to an LM-induced current (df = 7).

| Muscle | Mean (ms) | SD (ms) | T | P |
| --- | --- | --- | --- | --- |
| FDI | -0.01 | 0.68 | -0.1 | 0.960 |
| ADM | -0.44 | 1.68 | -0.7 | 0.486 |
| ECR | -0.88 | 1.22 | -2.1 | 0.079 |

FDI: first dorsal interosseous; ADM: abductor digiti minimi; ECR: extensor carpi radialis.

## 4. DISCUSSION

In this work, we assessed the effect of two different TMS-induced current directions (PA and LM) with respect to the individual brain morphology on the spatial representation of the MEP motor maps of 3 different muscles. We found that TMS-based motor mapping with PA-and LM-induced current directions reveal significantly different spatial representations of the same underlying motor area.

### 4.1 PA- and LM-induced currents identify different MEP motor maps

The effect of the TMS-induced current direction on the amplitude of the MEP has been described extensively[16][17]. Similar to these reports, we observed a significantly higher RMT for an LM-induced current with respect to PA-induced current.

In addition to a difference in RMT, we found that TMS-based motor mapping with a PA-induced current identifies a functional motor area located more anterior compared to an LM-induced current, for all the investigated muscles (FDI, ADM and ECR). This suggests that different current directions activate neuronal tissue in slightly different areas. It is possible that a PA-induced current has the propensity to activate corticospinal neurons in the premotor area rather than just in M1, which could explain the anterior displacement of the COG. This could occur due to the fact that currents perpendicular to the sulcal wall of the precentral gyrus more strongly activate pyramidal neurons in the cortical sheet as compared to currents flowing parallel to the sulcal wall[28]. Studies investigating direct cortical stimulation (DCS) and navigated TMS have shown that stimulation of the premotor cortex can indeed induce motor activity[11][29]. Unfortunately, both studies only used a PA-induced current direction during TMS motor mapping, which precluded a comparison with LM-induced current.

Besides the possibility that different TMS-induced current directions activate different anatomical areas, it is possible that different current directions activate different neuronal populations. Previous studies have found that LM-induced currents preferentially evoke activity in other cortical elements than PA currents. LM-induced currents have been shown to primarily elicit D-waves, which originate from axonal activation of cortico-spinal neurons (CSNs)[19]. PA-induced currents have been shown to preferentially evoke I-waves, which originate from synaptic inputs of the CSNs[20]. Consequently, PA-induced currents evoke MEPS with shorter latencies compared to LM-induced currents during stimulation of a single cortical target where the maximum MEP is evoked[27][30]. Our findings show that if all the MEPs in the MEP motor map are considered, the average latency of MEPs obtained with a PA-induced current is not significantly different from MEPs obtained with an LM-induced current. This suggests that the recruitment of different cortical elements depends on the direction is specific for the induced current with respect to the underlying morphology[27]. Therefore, it is unlikely that activation of different neuronal populations is the cause of the spatial shift between MEP motor maps obtained with PA- and LM-induced currents.

A more likely explanation comes from the interaction between the TMS-induced current direction and the local morphology of the stimulated cortex and the neuronal networks within the cortical layers. TMS evokes primary and secondary currents in the brain. The primary current is generated by the electrical field (E-field) resulting from the magnetic flux of the TMS coil, which also exists in a vacuum. The secondary currents arise from charge accumulations at tissue boundaries due to differences in tissue conductivity in different parts of the head (i.e. grey matter, white matter, skin)[31]. The secondary currents add up to the primary currents and can also contribute to MEP generation, which may therefore influence the spatial representation of the MEP motor map. It is possible that the common morphology of the precentral gyrus and the gradient in local conductivities affect secondary currents in such a way that PA-induced currents reveal an MEP motor area that is located anterior with respect to the motor area identified using LM-induced currents. Finite element modeling (FEM) has been proposed in order to make sensible claims about the exact effect of secondary currents on the difference between the spatial representations of the MEP motor areas obtained with different TMS induced current directions. This method can be used to model the TMS-induced currents based on electromagnetic principles, individual brain morphology and tissue conductivities[31]–[33]. In this way, Opitz et al. (2011) were able to illustrate the importance of local morphology by showing that the gyrus crowns are predominantly affected by TMS-induced currents rather than the sulcus area[34]. Further research is needed to validate these methods.

To the best of our knowledge, a similar experiment has been performed only once before by Wilson et al., who did not report a significant displacement of the motor area for different TMS-induced current directions in 5 participants[4]. However, they investigated different induced current directions (ML-induced current rather than LM-induced current) than in this study. Additionally, they delivered two stimuli with PA currents and two stimuli with ML currents for each grid point before moving to the next, without adjusting the stimulator output. The use of MRI-guided neuronavigation and real-time capturing of the TMS coil position allowed us to more accurately reproduce the TMS coil positions. These methodological differences do not allow a direct comparison with our results.

Clearly, further research is required to determine the mechanism(s) behind the spatial distribution of MEP-eliciting cortical targets in response to different TMS-induced current directions. This can be done using concurrent TMS-fMRI in order to directly measure TMS-induced brain activity, and/or FEM models, which use electromagnetic principles to determine TMS-induced activation patterns.

### 4.2 TMS-based motor mapping with LM-induced currents correspond better with fMRI

We found that the COG obtained with a PA induced current is located anterior to the location of the functional hand area as determined with fMRI. Although this separation was not significant for the FDI muscle (P = 0.05), this separation is in agreement with prior literature[12]–[15]. Interestingly, this separation was significant for the MEP motor map of the ADM muscle, which is expected as participants were instructed to perform movements involving the FDI instead of the ADM or ECR. The average separation between the TMS-based COG and the fMRI-based hand area was much smaller for an LM induced current compared to a PA induced current. This suggests that an MEP motor map obtained with an LM induced current corresponds better with the fMRI-based hand area.

The current golden standard for mapping of the motor area is pre- or intraoperative direct current stimulation (DCS), which has been compared to both fMRI and TMS localization techniques. Krieg et al. (2012) compared TMS-based motor mapping using a PA-induced current with fMRI-based motor mapping and DCS and found that motor maps obtained with TMS corresponded better with DCS compared to fMRI (average separations of 4.4 mm versus 9.8). They did not establish significance and they did not observe a unidirectional systematic deviation between these modalities. Forster et al. (2011) and Coburger et al. (2013) observed a similar trend[35][36]. These findings should be interpreted with caution as the findings are based on measurements in patients with brain tumors, as brain tumors have different tissue conductivities, potentially resulting in different TMS-induced current distributions.

Taken together, these results suggest that TMS-based motor maps obtained using a PA-induced current direction correspond better with the gold standard than fMRI-based maps, while motor maps obtained with an LM-induced current correspond better with fMRI-based maps. However, as both TMS-induced current directions have not been directly compared to DCS, we can only assume that motor maps obtained with a PA-induced current direction correspond best with DCS.

### 4.3 Navigated TMS-EMG can depict different muscle representations

A secondary aim of this study was to confirm the sensitivity of navigated TMS-EMG in the identification of separate MEP motor areas for different muscles: the FDI, ADM and ECR muscles. Wassermann et al. (1992) were the first to report that the maps of different muscles of the upper extremities showed a consistent organization within the precentral gyrus, as described by Penfield, based on TMS-EMG[37]. This has been observed by others as well[38]. However, 3Wassermann et al (1992) and Malcolm et al. (2006) did not quantify the separation between muscle motor maps.

We found that, at the group level, the COG of the MEP spatial distribution of the FDI muscle is significantly more lateral compared to the COG of the ADM muscle for both induced current directions, which is in line with layout of the motor homunculus as observed by Penfield[39]. Similar to our observations, Wilson et al. reported a significant separation between the centers of the FDI and ADM motor areas[4]. Although we found significantly separated COGs for the FDI and ADM muscles, the spatial distributions of the MEPs showed substantial overlap as well.

## 5. CONCLUSION

In conclusion, the spatial representations of the functional motor areas of 3 different muscles identified using navigated TMS-EMG depends on the direction of the TMS-induced current. TMS applied with induced currents perpendicular to the central sulcus (PA) identified a motor area that is located significantly more anterior with respect to an MEP motor area identified using currents induced parallel (LM) to the central sulcus. Furthermore, LM TMS-induced currents identified a motor map that more closely corresponds to the motor map identified using fMRI. The electrophysiological principles that underlie the difference between motor areas identified with different TMS-induced current directions remain unknown. We speculate that different TMS-induced current directions induce different distributions of secondary currents, which are shaped by cortical morphology and tissue conductivities. Further research is required to identify the mechanism(s) through which changes in the TMS-induced current direction affect the spatial distribution of motor-evoked potentials on brain surface. The current study clearly indicates that clinicians who are currently using navigated TMS-EMG to identify MEP motor maps for diagnostic or surgical planning procedures should carefully take into account the TMS-induced current direction in the interpretation and application of the TMS-based motor maps.

## Study funding/Acknowledgements

This work was supported by the DeNeCor project as part of the ENIAC Joint Undertaking, (ENIAC131003; http://eniac.eu). The funders had no role in study design, data collection and analysis, decision to publish, or preparation of the manuscript.

## REFERENCES

[1] A. T. Barker, R. Jalinous, and I. L. Freeston, “Non-invasive magnetic stimulation of human motor cortex,” Lancet, vol. 325, no. 8437, pp. 1106–1107, 1985.

[2] P. M. Rossini, D. Burke, R. Chen, L. G. Cohen, Z. Daskalakis, R. Di Iorio, V. Di Lazzaro, F. Ferreri, P. B. Fitzgerald, M. S. George, M. Hallett, J. P. Lefaucheur, B. Langguth, H. Matsumoto, C. Miniussi, M. A. Nitsche, A. Pascual-Leone, W. Paulus, S. Rossi, J. C. Rothwell, H. R. Siebner, Y. Ugawa, V. Walsh, and U. Ziemann, “Non-invasive electrical and magnetic stimulation of the brain, spinal cord, roots and peripheral nerves: Basic principles and procedures for routine clinical and research application: An updated report from an I.F.C.N. Committee,” Clin. Neurophysiol., vol. 126, no. 6, pp. 1071–1107, 2015.

[3] J. Lüdemann-Podubecká and D. A. Nowak, “Mapping cortical hand motor representation using TMS: A method to assess brain plasticity and a surrogate marker for recovery of function after stroke?,” Neurosci. Biobehav. Rev., vol. 69, pp. 239–251, 2016.

[4] S. a. Wilson, G. W. Thickbroom, and F. L. Mastaglia, “Transcranial magnetic stimulation mapping of the motor cortex in normal subjects,” J. Neurol. Sci., vol. 118, no. 2, pp. 134–144, 1993.

[5] P. E. Tarapore, M. C. Tate, A. M. Findlay, S. M. Honma, D. Mizuiri, M. S. Berger, and S. S. Nagarajan, “Preoperative multimodal motor mapping: a comparison of magnetoencephalography imaging, navigated transcranial magnetic stimulation, and direct cortical stimulation,” J. Neurosurg., vol. 117, no. 2, pp. 354–362, 2012.

[6] S. M. Krieg, J. Sabih, L. Bulubasova, T. Obermueller, C. Negwer, I. Janssen, E. Shiban, B. Meyer, and F. Ringel, “Preoperative motor mapping by navigated transcranial magnetic brain stimulation improves outcome for motor eloquent lesions,” Neuro. Oncol., vol. 16, no. 9, pp. 1274–1282, 2014.

[7] D. Frey, S. Schilt, V. Strack, A. Zdunczyk, J. R. Sler, B. Niraula, P. Vajkoczy, and T. Picht, “Navigated transcranial magnetic stimulation improves the treatment outcome in patients with brain tumors in motor eloquent locations,” Neuro. Oncol., vol. 16, no. 10, pp. 1365–1372, 2014.

[8] S. M. Krieg, N. Sollmann, T. Obermueller, J. Sabih, L. Bulubas, C. Negwer, T. Moser, D. Droese, T. Boeckh-Behrens, F. Ringel, and B. Meyer, “Changing the clinical course of glioma patients by preoperative motor mapping with navigated transcranial magnetic brain stimulation,” BMC Cancer, vol. 15, no. 1, pp. 1–11, 2015.

[9] N. Freundlieb, S. Philipp, A. Drabik, C. Gerloff, N. D. Forkert, and F. C. Hummel, “Ipsilesional motor area size correlates with functional recovery after stroke: A 6-month follow-up longitudinal TMS motor mapping study,” Restor. Neurol. Neurosci., vol. 33, no. 2, pp. 221–231, 2015.

[10] M. Yarossi, S. Adamovich, and E. Tunik, “Sensorimotor cortex reorganization in subacute and chronic stroke: A neuronavigated TMS study,” Conf. Proc Annu. Int. Conf. IEEE Eng. Med. Biol. Soc. IEEE Eng. Med. Biol. Soc. Annu. Conf., vol. 2014, pp. 5788–5791, 2014.

[11] T. Moser, L. Bulubas, J. Sabih, N. Sollmann, B. Meyer, F. Ringel, and S. M. Krieg, “Resection of Navigated Transcranial Magnetic Stimulation-Positive Prerolandic Motor Areas Causes Permanent Impairment of Motor Function,” vol. 81, no. 1, pp. 99–110, 2018.

[12] R. Sparing, D. Buelte, I. G. Meister, T. Pauš, and G. R. Fink, “Transcranial magnetic stimulation and the challenge of coil placement: A comparison of conventional and stereotaxic neuronavigational strategies,” Hum. Brain Mapp., vol. 29, no. 1, pp. 82–96, 2008.

[13] U. Herwig, K. Kölbel, A. P. Wunderlich, A. Thielscher, C. von Tiesenhausen, M. Spitzer, and C. Schönfeldt-Lecuona, “Spatial congruence of neuronavigated transcranial magnetic stimulation and functional neuroimaging,” Clin. Neurophysiol., vol. 113, no. 4, pp. 462–468, 2002.

[14] M. Lotze, R. J. Kaethner, M. Erb, L. G. Cohen, W. Grodd, and H. Topka, “Comparison of representational maps using functional magnetic resonance imaging and transcranial magnetic stimulation,” Clin. Neurophysiol., vol. 114, no. 2, pp. 306–312, 2003.

[15] S. F. W. Neggers, T. R. Langerak, D. Schutter, R. C. W. Mandl, N. F. Ramsey, P. J. J. Lemmens, and A. Postma, “A stereotactic method for image-guided transcranial magnetic stimulation validated with fMRI and motor-evoked potentials,” Neuroimage, vol. 21, no. 4, pp. 1805–1817, 2004.

[16] K. R. Mills, S. J. Boniface, and M. Schubert, “Magnetic brain stimulation with a double coil: the importance of coil orientation,” Electroencephalogr. Clin. Neurophysiol. Potentials Sect., vol. 85, no. 1, pp. 17–21, 1992.

[17] L. Richter, G. Neumann, S. Oung, A. Schweikard, and P. Trillenberg, “Optimal Coil Orientation for Transcranial Magnetic Stimulation,” PLoS One, vol. 8, no. 4, pp. 1–10, 2013.

[18] T. Kammer, M. Vorwerg, and B. Herrnberger, “Anisotropy in the visual cortex investigated by neuronavigated transcranial magnetic stimulation,” Neuroimage, vol. 36, no. 2, pp. 313–321, 2007.

[19] L. J. Volz, M. Hamada, J. C. Rothwell, and C. Grefkes, “What Makes the Muscle Twitch: Motor System Connectivity and TMS-Induced Activity,” Cereb. Cortex, vol. 25, no. 9, pp. 2346–2353, 2015.

[20] V. Di Lazzaro, P. Profice, F. Ranieri, F. Capone, M. Dileone, A. Oliviero, and F. Pilato, “I-wave origin and modulation,” Brain Stimul., vol. 5, no. 4, pp. 512–525, 2012.

[21] V. Di Lazzaro, U. Ziemann, and R. N. Lemon, “State of the art: Physiology of transcranial motor cortex stimulation,” Brain Stimul., vol. 1, no. 4, pp. 345–362, 2008.

[22] M. Lotze, M. Erb, H. Flor, E. Huelsmann, B. Godde, and W. Grodd, “fMRI evaluation of somatotopic representation in human primary motor cortex,” Neuroimage, vol. 11, no. 5 I, pp. 473–481, 2000.

[23] S. Rossi, M. Hallett, P. M. Rossini, A. Pascual-Leone, G. Avanzini, S. Bestmann, A. Berardelli, C. Brewer, T. Canli, R. Cantello, R. Chen, J. Classen, M. Demitrack, V. Di Lazzaro, C. M. Epstein, M. S. George, F. Fregni, R. Ilmoniemi, R. Jalinous, B. Karp, J. P. Lefaucheur, S. Lisanby, S. Meunier, C. Miniussi, P. Miranda, F. Padberg, W. Paulus, A. Peterchev, C. Porteri, M. Provost, A. Quartarone, A. Rotenberg, J. Rothwell, J. Ruohonen, H. Siebner, G. Thut, J. Valls-Sol??, V. Walsh, Y. Ugawa, A. Zangen, and U. Ziemann, “Safety, ethical considerations, and application guidelines for the use of transcranial magnetic stimulation in clinical practice and research,” Clin. Neurophysiol., vol. 120, no. 12, pp. 2008–2039, 2009.

[24] W. D. Penny, K. J. Friston, J. T. Ashburner, S. J. Kiebel, and T. E. Nichols, Statistical parametric mapping: the analysis of functional brain images. Academic press, 2011.

[25] J. Ashburner and K. J. Friston, “Unified segmentation,” Neuroimage, vol. 26, no. 3, pp. 839–851, 2005.

[26] I. Laakso, A. Hirata, and Y. Ugawa, “Effects of coil orientation on the electric field induced by TMS over the hand motor area,” Phys. Med. Biol., vol. 59, no. 1, pp. 203–218, 2014.

[27] M. Hamada, N. Murase, A. Hasan, M. Balaratnam, and J. C. Rothwell, “The role of interneuron networks in driving human motor cortical plasticity,” Cereb. Cortex, vol. 23, no. 7, pp. 1593–1605, 2013.

[28] T. Kammer, S. Beck, A. Thielscher, U. Laubis-Hermann, and H. Topka, “Motor threshold in humans: a transcranial magnetic stimulation study comparing different pulse waveforms, current directions and stimulator types,” Clin. Neurophysiol., vol. 112, pp. 250–8, 2001.

[29] L. Bulubas, J. Sabih, A. Wohlschlaeger, N. Sollmann, T. Hauck, S. Ille, F. Ringel, B. Meyer, and S. M. Krieg, “Motor areas of the frontal cortex in patients with motor eloquent brain lesions,” J. Neurosurg., vol. 125, no. 6, pp. 1431–1442, 2016.

[30] V. Di Lazzaro, F. Pilato, M. Dileone, P. Profice, F. Capone, F. Ranieri, G. Musumeci, A. Cianfoni, P. Pasqualetti, and P. A. Tonali, “Modulating cortical excitability in acute stroke: a repetitive TMS study,” Clin. Neurophysiol., vol. 119, no. 3, pp. 715–723, 2008.

[31] S. F. W. Neggers, P. I. Petrov, S. Mandija, I. E. C. Sommer, and N. A. T. van den Berg, Understanding the biophysical effects of transcranial magnetic stimulation on brain tissue: The bridge between brain stimulation and cognition., 1st ed., vol. 222. Elsevier B.V., 2015.

[32] A. Thielscher, A. Antunes, and G. B. Saturnino, “Field modeling for transcranial magnetic stimulation: A useful tool to understand the physiological effects of TMS?,” in Engineering in Medicine and Biology Society (EMBC), 2015 37th Annual International Conference of the IEEE, 2015, pp. 222–225.

[33] A. Bungert, A. Antunes, S. Espenhahn, and A. Thielscher, “Where does TMS Stimulate the Motor Cortex? Combining Electrophysiological Measurements and Realistic Field Estimates to Reveal the Affected Cortex Position,” Cereb. Cortex, vol. 27, no. 11, pp. 5083–5094, 2017.

[34] A. Opitz, M. Windhoff, R. M. Heidemann, R. Turner, and A. Thielscher, “How the brain tissue shapes the electric field induced by transcranial magnetic stimulation,” Neuroimage, vol. 58, no. 3, pp. 849–859, 2011.

[35] M. T. Forster, E. Hattingen, C. Senft, T. Gasser, V. Seifert, and A. Szelényi, “Navigated transcranial magnetic stimulation and functional magnetic resonance imaging: Advanced adjuncts in preoperative planning for central region tumors,” Neurosurgery, vol. 68, no. 5, pp. 1317–1324, 2011.

[36] J. Coburger, C. Musahl, H. Henkes, D. Horvath-Rizea, M. Bittl, C. Weissbach, and N. Hopf, “Comparison of navigated transcranial magnetic stimulation and functional magnetic resonance imaging for preoperative mapping in rolandic tumor surgery,” Neurosurg. Rev., vol. 36, no. 1, pp. 65–75, 2013.

[37] E. M. Wassermann, L. M. McShane, M. Hallett, and L. G. Cohen, “Noninvasive mapping of muscle representations in human motor cortex,” Electroencephalogr. Clin. Neurophysiol. Evoked Potentials, vol. 85, no. 1, pp. 1–8, 1992.

[38] M. P. Malcolm, W. J. Triggs, K. E. Light, O. Shechtman, G. Khandekar, and L. J. G. Rothi, “Reliability of motor cortex transcranial magnetic stimulation in four muscle representations,” Clin. Neurophysiol., vol. 117, no. 5, pp. 1037–1046, 2006.

[39] W. Penfield and T. Rasmussen, “The cerebral cortex of man; a clinical study of localization of function.,” 1950.

